# Variations of a theme - crystal forms of the amino acid transporter MhsT

**DOI:** 10.1101/2022.05.03.490450

**Authors:** Caroline Neumann, Dorota Focht, Sofia Trampari, Joseph Lyons, Poul Nissen

## Abstract

The bacterial amino acid transporter MhsT of the SLC6A family was crystallized in complex with different substrates in order to understand the determinants of substrate specificity of the transporter. Surprisingly, crystals of the different MhsT-substrate complexes showed interrelated, but different crystal packing arrangements. Space group assignment and structure determination of these different crystal forms presented challenging combinations of pseudosymmetry, twinning and tNCS.

**Synopsis:** An unusual case of protein-substrate complexes obtained in similar conditions, but containing different packing arrangements. The crystals exhibit a combination of various crystal imperfections (pseudosymmetry, twinning and tNCS) masking the true crystal symmetries and challenging data processing and structure determination.

## 1. Introduction

Macromolecular crystals often suffer from imperfections that cause difficulties in space group determination, data processing and refinement. Twinning is one of the most often encountered crystal defects. Different types of twinning are known: Epitaxial or non-merohedral twinning is present, when lattices of the twin domains overlap in fewer than three dimensions, therefore making the diffraction patterns look abnormal. This kind of twinning can be detected by visual inspection of the diffraction images, and in some cases the diffraction spots belonging to the individual domains can be identified and processed separately (Liang *et al*., 1996, Lietzke *et al*., 1996). In contrast, merohedral twinning is characterized by a complete overlap of real space lattices from the twin domains resulting in superposition of the reciprocal lattice, hence appearing normal (Yeates, 1997).

Merohedral twinning is detected by intensity distribution analyses, which will deviate from theoretical Wilson statistics due to averaging of independent lattices that reduces the variation in intensity distributions (Wilson, 1949, Chandra *et al*., 1999, Stanley, 1972). In merohedral twinning the holohedry belongs to a higher point group than the symmetry of the Laue class and therefore, coset decomposition of the holohedry with regards to the Laue class is a method to determine the possible twin laws (Flack, 1987). These twin laws can be used to “detwin” the crystal and calculate the true intensities in cases of twin fractions much smaller than 0.5, or to refine twinned crystal structures with use of the twin law in the cases of perfect twinning (Yeates, 1997).

Merohedral twinning is only present in point groups belonging to crystal systems containing several Laue classes, such as point groups 3, 4, 6, 23 and 32 (hexagonal setting) (Yeates, 1997). Therefore, for point group 2, merohedral twinning is generally not possible, but there are exceptions. For example, in the fortuitous case of *β* ≈ 90° an orthorhombic unit cell can be mimicked and twinning becomes possible (Larsen *et al*., 2002). Here, the holohedry exhibits *mmm* point group symmetry, whereas the crystal structure has point group 2, which can be caused by two possible equivalent twin laws along **a** (h,-k,-l) or **c** (-h,-k,l). This kind of twinning is called pseudomerohedral twinning with the orthorhombic and monoclinic point groups belonging to two different crystal systems (Parsons, 2003). In contrast to merohedral twinning, the lattices of the different twin domains overlap only approximately in three dimensions and therefore, the spots in the diffraction pattern will not superpose completely (Yeates, 1997), often appearing as streaky reflections.

Pseudosymmetry is another phenomenon that may mask the true crystal symmetry. It is often observed, where non-crystallographic symmetry (NCS) operators lie close to crystallographic operators, and like in the case of twinning, the holohedry has a higher point group than the crystal. A typical hallmark of pseudosymmetry are problems related to high R-factors during refinement (Zwart *et al*., 2008).

A third common crystal phenomenon that impacts data processing and structure refinement is translational NCS (tNCS). Here, the NCS related molecules are related only by a translation, while the orientation stays almost the same. This brings in a modulation of the diffraction pattern by existence of systematic weak and strong spots arising from the fact that the related molecules contribute similar structure factor amplitudes, but different phases (Read *et al*., 2013). Translational NCS can be detected in the Patterson map by the presence of non-origin peaks with a height of at least 20% of the origin peak. The smaller the difference in orientation between the tNCS related molecules, the more significant will the effect of tNCS be on data processing, phasing and refinement.

In this report, we describe three different crystal forms of the multihydrophobic amino acid transporter (MhsT) with complications of pseudosymmetry, different degrees of pseudomerohedral twinning, and translational NCS. MhsT is an amino acid transporter from *Bacillus halodurans* that transports a variety of hydrophobic L-amino acids. It is an orthologue of the mammalian neurotransmitter:sodium symporters and amino acid transporters of the SLC6 family. MhsT substrates range from the bulky, aromatic substrates: tryptophan (Trp), tyrosine (Tyr) and phenylalanine (Phe) to the smaller, branched aliphatic amino acids: valine (Val), leucine (Leu) and isoleucine (Ile) (Quick & Javitch, 2007). The initial aim of our study was to understand determinants of substrate specificity of MhsT through co-crystallization with all substrates. Six different structures were determined and together with the previously published MhsT-Trp complex (Malinauskaite *et al*., 2014), a substrate recognition mechanism involving the unwound part of TM6 was elucidated (Focht *et al*., 2021). Somewhat surprisingly, however, we observed that different crystal forms emerged, yet displaying related intermolecular crystal packing arrangements. MhsT in complex with 4-F-Phe, Tyr, Phe and the previously determined Trp, crystallize in a P2 space group with the long axis along **c**. In the case of the smaller ligands (Val, Leu) a slight change in packing occurs and the space group changes to P2_1_ with new parameters: a_P21_= a_P2_, b_P21_=2c_P2_, c_P21_= b_P2_ and *β* ≈ 90°. The unit cell in the P2_1_ crystal form is approximately twice the volume of the P2 form and accommodates two MhsT complexes instead of one in the asymmetric unit, related by rotational NCS. The P2 form was never observed for the smaller aliphatic substrates, but the P2_1_ crystal form on the other hand was also observed for the aromatic substrates, although higher quality datasets were obtained in P2.

Another crystal packing variation occurred in the case of the MhsT-Ile complex, which crystallized in a different P2_1_ crystal form, now with unit cell parameters: a_P21_= a_P2_, b_P21_=2b_P2_ and c_P21_= c_P2_. Again, the unit cell is twice as large as for the P2 form with the asymmetric unit containing two MhsT molecules, however now related by translational NCS. Table 1 presents an overview of the different complexes, with a description of the space group, unit cell parameters and data statistics.

**Table 1.**
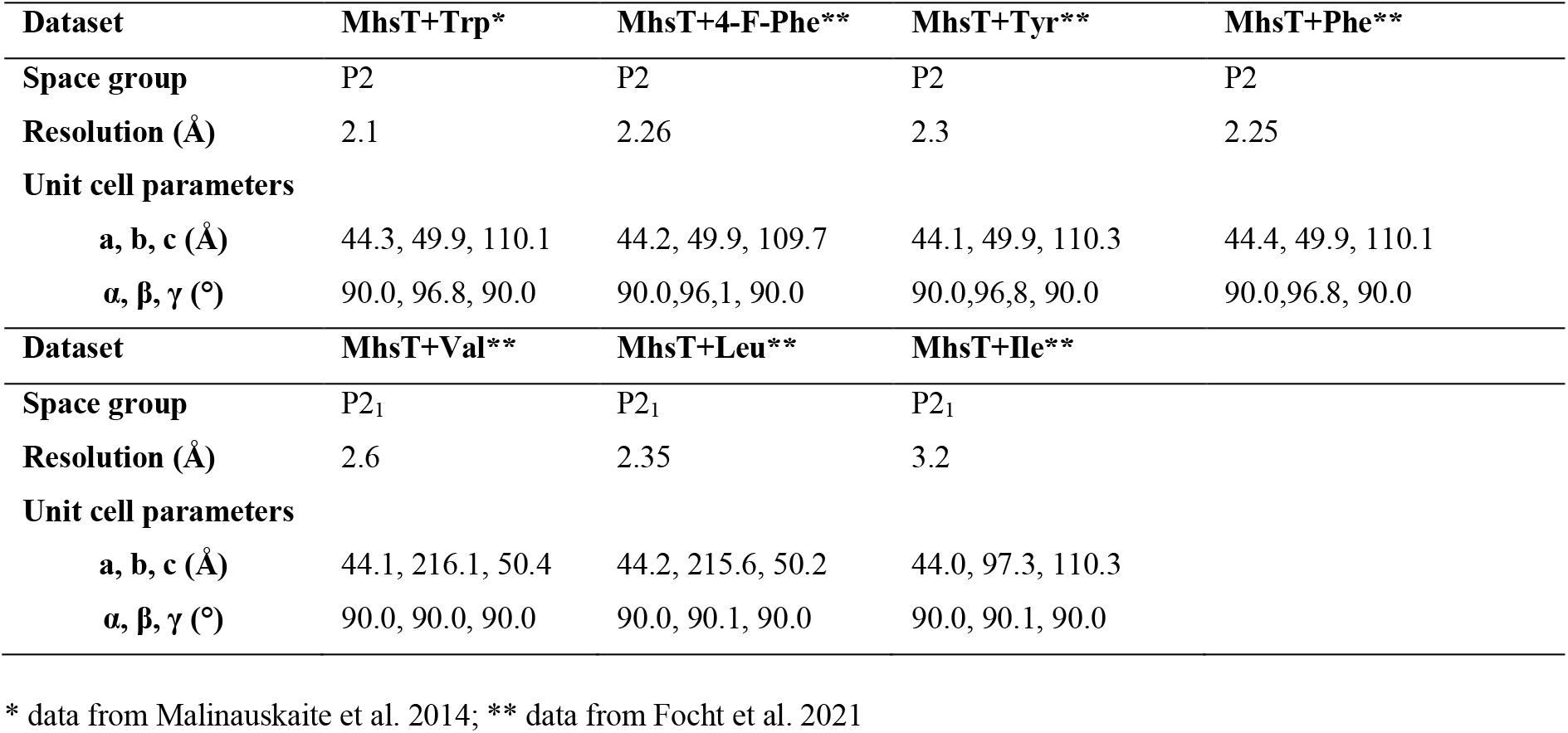
Space group and unit cell parameters of the different MhsT complexes.

While data processing and refinement were straightforward in the case of structures determined in P2, the P2_1_ cases turned out to be more challenging, especially because of the presence of pseudosymmetry and twinning in the Val- and Leu-bound complexes, and translational NCS in the case of MhsT-Ile, obscuring space group assignment and refinement.

Here we describe how the crystal imperfections were identified and accounted for to ensure a proper refinement of molecular models.

## 2. Methods

### 2.1. Protein expression, purification and crystallization

Expression, purification and crystallization of MhsT in complex with its different substrates were described previously (Focht *et al*., 2021), based on earlier studies (Malinauskaite *et al*., 2014, Quick & Javitch, 2007). In short, all crystal forms were obtained in a 0.3-0.5 M NaCl containing buffer at pH 7.0, 5% Trimethylamine N-oxide (TMANO) and using 14-24% PEG400 as the precipitant at 19°C. Obtained crystals were harvested at 4°C.

### 2.2. Crystal morphology and data collection

Crystals of MhsT in complex with aromatic substrates (Phe, 4-F-Phe, Tyr) and branched, aliphatic substrates (Val or Leu) were small, three-dimensional rodlike crystals (Figure 1A), with lengths ranging from 30 to 50 µm similar to MhsT-Trp crystals (Malinauskaite *et al*., 2014). Data collection was performed at Diamond Light Source at beamlines I24 and I04 (Focht *et al*., 2021).

**Figure 1.**
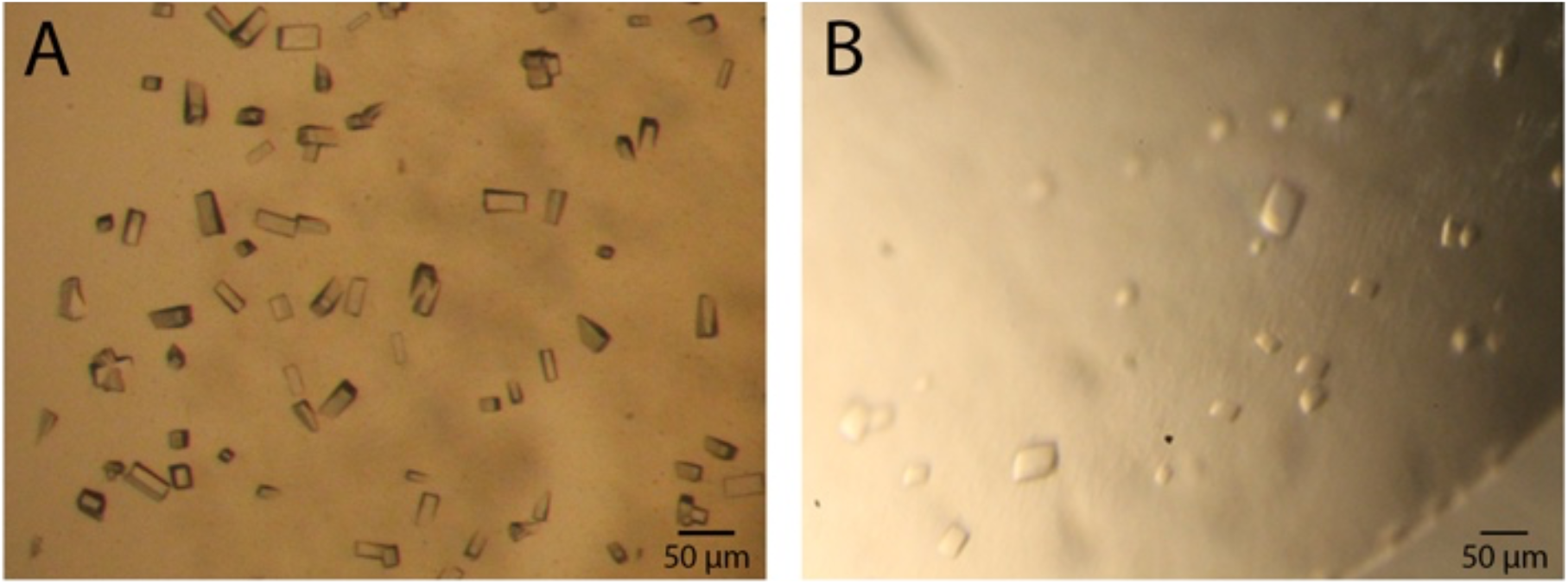
Crystals of MhsT. A) P21 crystal form of MhsT-Val exhibiting P2221 pseudosymmetry and negligible twinning, and B) similar P21 crystal form of MhsT-Leu with twin fraction 0.43. C) MhsT-Ile crystallizes in a different P21 form with translational non-crystallographic symmetry

The exception were the crystals of MhsT-Ile that appeared as flat plates with dimensions up to 70 *µ*m and about 5-10 *µ*m thick (Figure 1B). The datasets were collected at the Swiss Light Source at beamline PXI. The MhsT-Ile crystals were very sensitive to radiation damage and complete datasets could not be obtained from single crystals (Focht *et al*., 2021).

## 3. Two datasets with pseudosymmetry and pseudomerohedral twinning

### 3.1. Data processing of MhsT-Val and MhsT-Leu datasets

The MhsT-Val and MhsT-Leu datasets were initially processed by use of the XDS package (Kabsch, 2010) and CCP4 program (Matthews, 1968, Winn *et al*., 2011) in the orthorhombic P222_1_ space group with systematic absences along **c**. The processing resulted in an overall R_merge_ of 9.9% in the case of MhsT-Val and 12.7% in the case of MhsT-Leu, indicating seemingly acceptable merging statistics and space group assignment. Initial phases were obtained by use of molecular replacement in *Phenix*.*Phaser-MR* using Mhst-Trp (PDB 4US3) without TM5 and ligands as a search model and identifying one molecule in the asymmetric unit (Matthew coefficient of 2.57, solvent content of 52.1% for MhsT-Val, and Matthew coefficient of 2.55, solvent content of 51.9% for MhsT-Leu). The molecular replacement solutions were clear in both cases (PAK=0, LLG=822 and TFZ= 18.2 for MhsT-Val and PAK=0, LLG= 1262 and TFZ= 19.1 for MhsT-Leu), However, refinement stalled in both cases at R_work_/R_free_ of about 0.38/0.43.

These R-factor statistics indicated that the structure did not explain the diffraction data well, and that the assignment of P222_1_ space group symmetry was potentially incorrect. The presence of pseudosymmetry and/or twinning in the datasets was suspected to make the diffraction pattern resemble an orthorhombic space group due to a fortuitous value of *β* ≈ 90°. Therefore, the three monoclinic “Translationengleiche” subgroups of P222_1_ were explored to investigate which twofold or screw operator present in the orthorhombic space group remained valid (if any) as a crystallographic operator in a monoclinic space group.

In the case of MhsT-Val, the merging R-factors for two P2 space groups were markedly increased (Table 2), suggesting that they too were not valid space groups for this dataset, and indeed, they resulted also in high model R-factors in refinement. However, processing in the P2_1_ space group with the long axis (the c axis in P222_1_) now along **b**, merged with proper statistics and an overall R_merge_=0.109. Molecular replacement located two molecules in the asymmetric unit (Matthews coefficient of 2.57, solvent content of 52.1%), yielding a single solution with PAK=0, LLG=5863.578 and TFZ= 53.9. The model refinement proceeded smoothly and converged with R_work_/R_free_ at 0.207/0.237 with high quality electron density maps also for unmodeled parts.

**Table 2.**
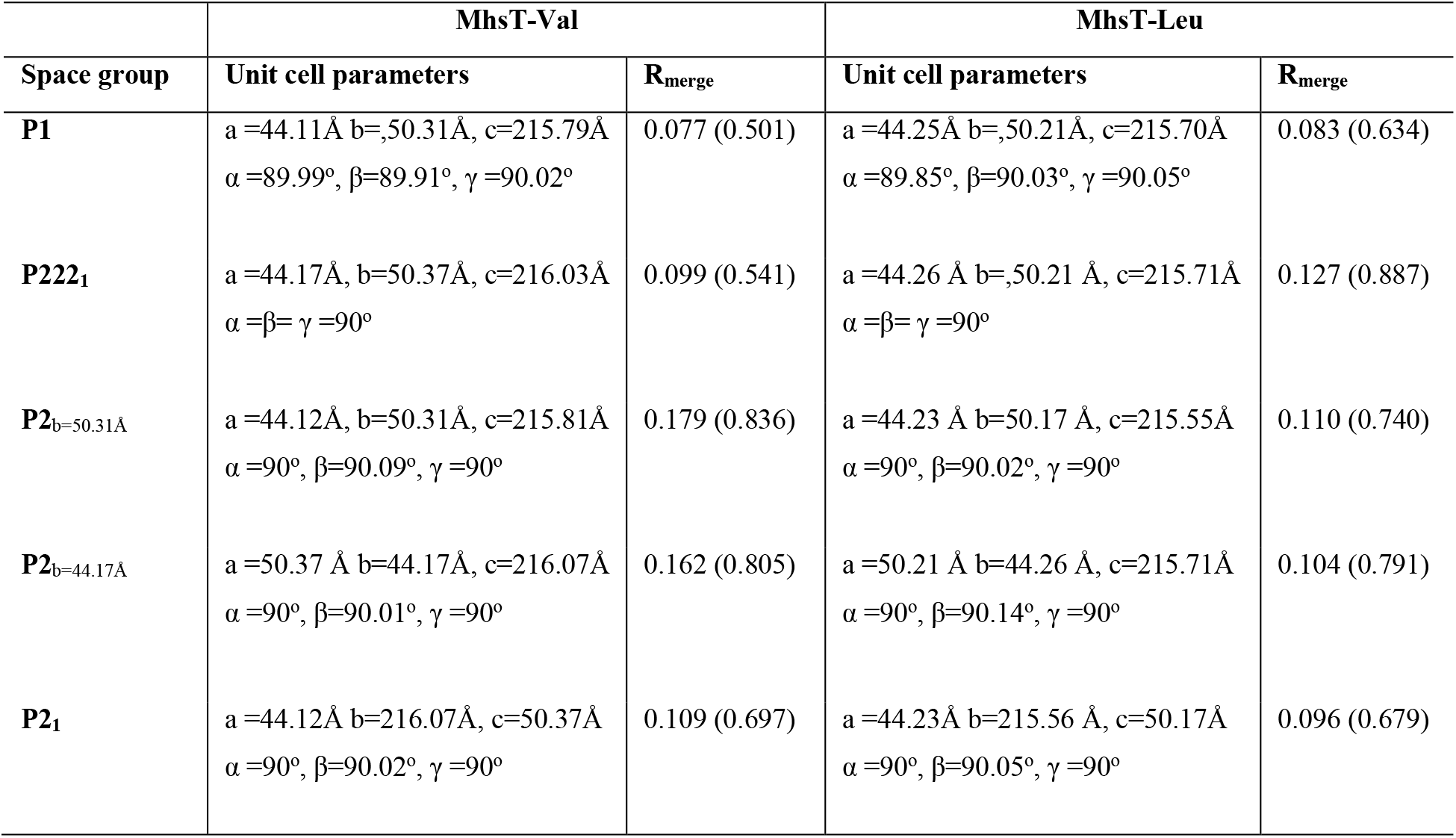
Processing of MhsT-Val and MhsT-Leu in different space groups.

Similarly, parallel runs of molecular replacement and initial refinement were performed for MhsT-Leu yielding initial R_work_/R_free_ values of 0.26/0.31 for P2_1,_ and 0.35/0.42 for P2_b=44.26Å_, and 0.31/0.37 for P2_b=50.17Å_. These results clearly indicated that the P2_1_ space group again was the correct assignment, similar to MhsT-Val. The complete processing and refinement statistics for the two datasets processed in P2_1_ can be seen in Table 3.

**Table 3.** Processing and refinement statistics for MhsT-Val and MhsT-Leu (Focht *et al*., 2021)

| Data collection | MhsT-Val | MhsT-Leu | MhsT-Ile |
| --- | --- | --- | --- |
| **Beamline** | DLS-I02 | DLS-I04 | SLS-PXI |
| **Space group** | P2<sub>1</sub> | P2<sub>1</sub> | P2<sub>1</sub> |
| **Unit cell parameters** |  |  |  |
| a, b, c (Å) | 44.1, 216.1, 50.4 | 44.2, 215.6, 50.2 | 44.0, 97.3, 110.9 |
| $\alpha, \beta, \gamma$ (°) | 90, 90.02, 90 | 90, 90.05, 90 | 90.0, 96.14, 90.0 |
| **Resolution range (Å)** | 49-2.60<br>(2.72-2.60) | 45-2.35<br>(2.43-2.35) | 43.7 – 3.10<br>(3.31-3.10) |
| **Number of reflections total/unique** | 97704/28703 | 114897/38300 | 37578/13889 |
| **Wilson B-factor** | 49.3 | 37.4 | 63.0 |
| **Completeness** | 99.5 (99.7) | 98.5 (98.5) | 81.6 (83.8) |
| **R<sub>merge</sub>** | 0.117 (0.950) | 0.096 (0.679) | 0.195 (0.815) |
| **R<sub>pim</sub>** | 0.075 (0.607) | 0.066 (0.502) | 0.124 (0.532) |
| **Mean I/sd (I)** | 8.0 (1.2) | 8.1 (1.4) | 3.6 (1.2) |
| **Redundancy** | 3.4 (3.4) | 3.0 (2.6) | 2.7 (2.5) |
| **CC1/2** | 0.994 (0.489) | 0.996 (0.511) | 0.989 (0.636) |
| **Twin fraction derived** | 0.065 | 0.443 | N/A |
| **Refinement** |  |  |  |
| **Twin law** | not used | h, -k, -l | N/A |
| **R<sub>work</sub>/R<sub>free</sub>** | 0.207/0.237 | 0.185/0.222 | 0.277/0.305 |
| **No. molecules/a.s.u.** | 2 | 2 | 2 |
| **Number of atoms** |  |  |  |
| Protein | 6674 | 6678 | 6630 |
| Ligand | 16 | 18 | 18 |
| Ions | 4 | 4 | 4 |
| Detergent/lipid | 357 | 176 | 47 |
| Water | 73 | 46 | 19 |
| **Average B- factor (Å<sup>2</sup>)** |  |  |  |
| Protein | 56.9 | 41.4 | 56.7 |
| Ligand | 48.8 | 34.1 | 52.6 |
| Ions | 49.6 | 33.3 | 52.8 |
| Detergent/lipid | 68.4 | 51.7 | 58.2 |
| Water | 55.4 | 37.9 | 52.2 |
| **RMS** |  |  |  |
| Bond lengths (Å) | 0.003 | 0.002 | 0.002 |
| Bond angles (°) | 0.595 | 0.505 | 0.578 |
| **Ramachandran (%)** |  |  |  |
| Favourable | 96.4 | 96.9 | 95.7 |
| Outliers | 0 | 0 | 0.23 |

a) Values in parentheses are for highest resolution shell.
b) MhsT-Leu was refined against the h,-k,-l twin law, whereas MhsT-Val was refined without the twin law as it did not have any significant influence on the R-factors during refinement.
c) 5% of the datasets was chosen for the R<sub>free</sub> Sets, additionally in the cases with 2 molecules in the asymmetric unit, R<sub>free</sub> flag was assigned in thin resolution shells.

However, we noted that for MhsT-Leu, the data scaling statistics looked comparable in all three monoclinic assignments, i.e., with the two P2 space groups having only a slightly increased R_merge_ compared to P2_1_. This seemingly different behavior of the MhsT-Val and-Leu data was investigated further and explained in the section further below.

### 3.2. Pseudosymmetry of the MhsT-Val and MhsT-Leu crystals

The presence and orientation of crystallographic and non-crystallographic rotational and screw axes in the two datasets were analysed by use of the Patterson self-rotation function (Rossmann & Blow, 1962). Figure 2 presents a stereographic projection of the self-rotation function at *k* = 180°, indicating 222 symmetry or 222 pseudosymmetry with the NCS axes lying close to crystallographic axes, which therefore candidates also as a possible twin axis.

**Figure 2.**
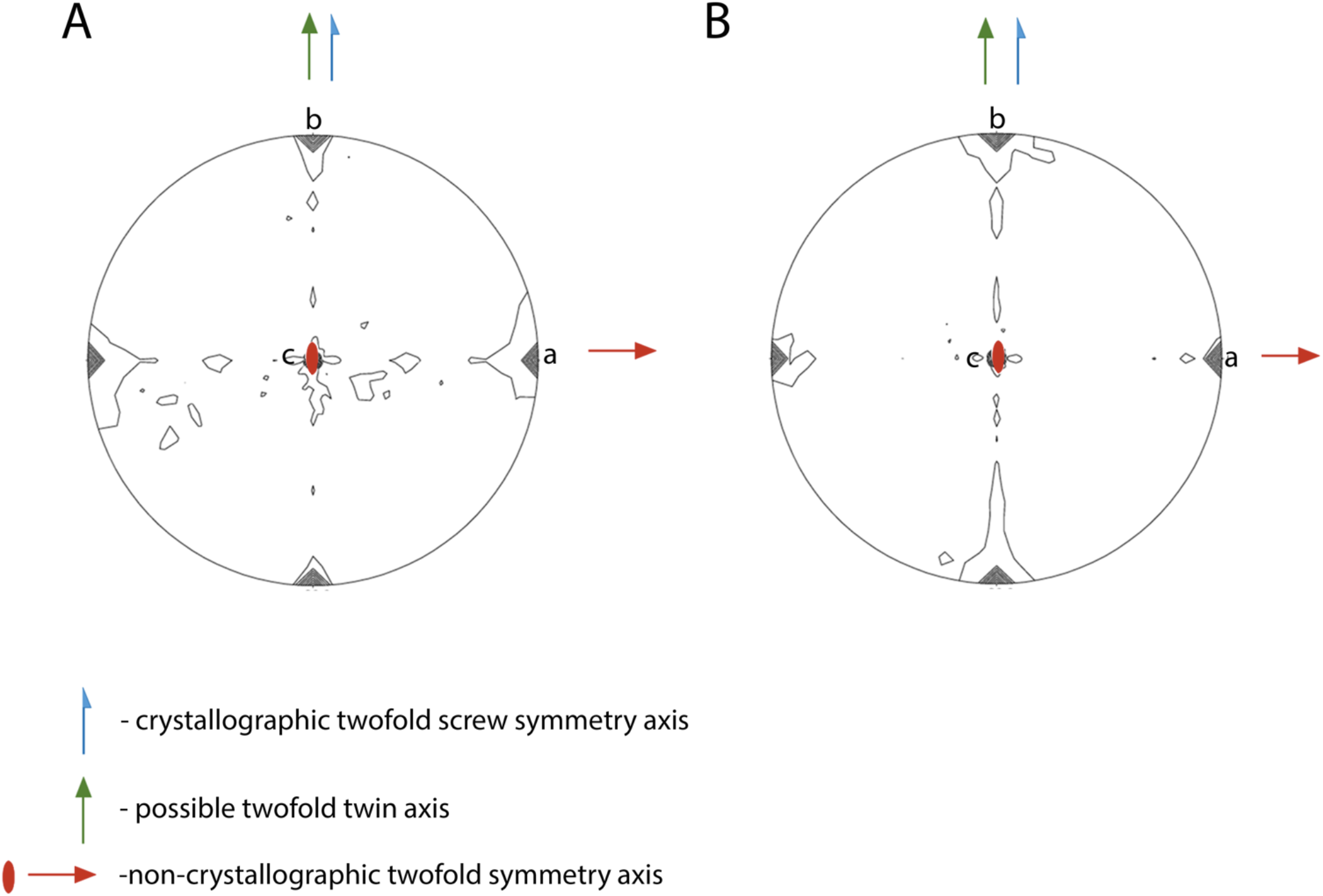
Self-rotation function of A) MhsT-Val and b) MhsT-Leu at κ=180° section. Low resolution limit=7Å, high resolution limit=3Å, radius of integration= 32Å. The crystallographic twofold screw symmetry axis is present along B), the non-crystallographic twofold symmetry axis and twin axis is present along a and the third twofold axis is present along c.

The P222_1_ pseudosymmetry prompted us to investigate the NCS operators with *Find NCS operators* in *Phenix*. The following operators were found for the two datasets:

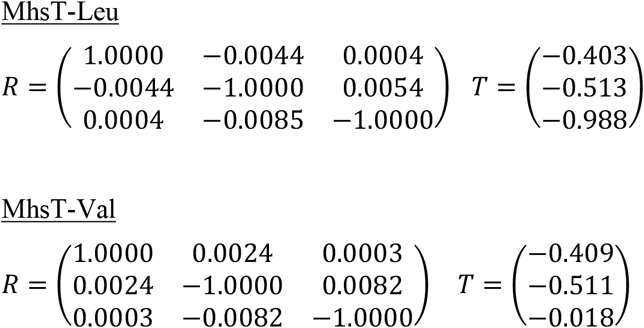

Hence, the NCS operator approximates a crystallographic twofold axis along **a**, which together with the crystallographic twofold axis along **b** generates a third twofold operator parallel to **c** and a P222_1_ pseudosymmetry. This operator is assumed to be crystallographic, when processing is done in the orthorhombic space group. However, because of the small imperfections of the NCS operator and translations combined, the refinement in the orthorhombic space group stalls at high model R-factors as indicated above.

### 3.3. Pseudomerohedral twinning of MhsT-Leu

Intensity analyses were performed on the monoclinic datasets in *Phenix*.*Xtriage* (Zwart *et al*., 2005) as *β* approaches 90° for both MhsT-Val and -Leu, making pseudomerohedral twinning possible. The Wilson ratio and secondary intensity moments (Stanley, 1972, Rees, 1980) of centric and acentric reflections hinted at twinning in both datasets (data not shown) with the h,-k,-l twin law. Additionally, the more robust local intensity difference analysis of the datasets was performed by use of “The Merohedral Twinning Server” (https://services.mbi.ucla.edu/Twinning/) applying the Padilla-Yeates Algorithm (Padilla & Yeates, 2003). Presence of twinning is indicated in both datasets (Figure 3), but with a markedly higher degree of twinning in the case of MhsT-Leu.

**Figure 3.**
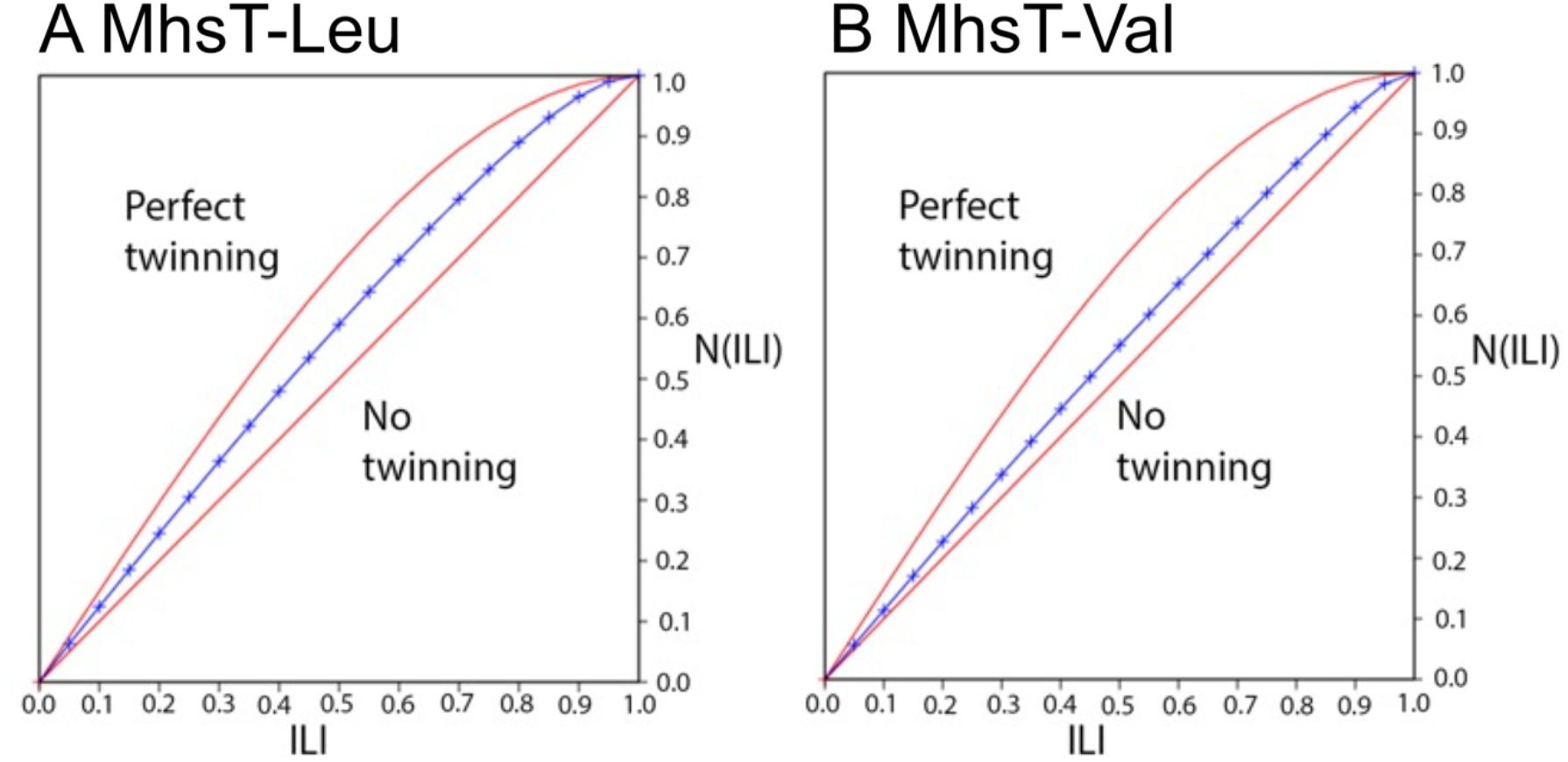
Graphs presenting statistics in the A) MhsT-Leu and B) MhsT-Val datasets with approx. 44% twinning and negligible twinning present

The presence of a two-fold NCS operator along **a** can bias twinning detection to indicate a twin operator along **a**, because the scaling statistics analysis assumes that the related reflections are otherwise independent (Yeates & Rees, 1987, Lebedev *et al*., 2006), which already by the NCS operator they clearly are not. At the same time, in such cases where the NCS operator coincides with a crystal axis make twinning highly probable (Lebedev *et al*., 2006). Generally, twin fractions can be determined in several ways (Britton, 1972, Rees, 1980, Fisher & Sweet, 1980, Murray-Rust, 1973, Yeates, 1997), however many of these tests do not give an accurate estimate of the twin fraction in cases like this. Therefore, a maximum likelihood test that takes the NCS axis under consideration was used in *Phenix*.*Xtriage*. The obtained twin fraction for MhsT-Val was 0.065, while it was 0.443 for MhsT-Leu. In comparison, the Britton test (Britton, 1972) gave twin fractions of 0.173 in the case of MhsT-Val and 0.447 for MhsT-Leu.

A significant drop (e.g. 3-10%) in model R-factors is to be expected in cases with high degrees of twinning, when comparing refinement with and without twin law. Indeed, for the MhsT-Leu dataset, structure refinement with the h,-k,-l twin law resulted in a drop of refinement R-factors from R_work_/R_free_ of 0.266/0.298 to R_work_/R_free_ of 0.185/0.222. However, in the MhsT-Val complex with a low degree of twinning, the R_work_/R_free_ was 0.207/0.237 without the use of the twin law and 0.194/0.223 when the h,-k,-l twin law was used; i.e., an almost negligible difference. As refinement with the twin law also did not improve the electron density maps for MhsT-Val, we refined the MhsT-Val structure without use of the twin law.

The significant difference in twin fractions observed between the two datasets explains the low merging factors of the MhsT-Leu dataset in all three monoclinic subgroups of P222_1_. In the case of a dataset belonging to the P222_1_, proper scaling and low merging factors would be expected in its type I maximal non-isomorphous subgroups. Therefore, in a case of two monoclinic datasets containing twin fractions of ∼0 and ∼0.443 it could be expected that the dataset with almost perfect twinning would scale better in all three subgroups, as the diffraction pattern approximates an orthorhombic setting more than the dataset with the lower twin fraction. However, in the case of pseudosymmetry present at the same time, like here, the merging statistics will also appear valid in the case of P222_1_, even though the twin fraction is small or non-existent (Parsons, 2003).

### 3.4. Crystal packing explains the pseudosymmetry of MhsT-Val and MhsT-Leu

In order to visualize the pseudosymmetry in the crystal structure, a closer analysis of the crystal packing was made. The two molecules in the P2_1_ asymmetric unit, molecules A and B (Figure 4A and 4B), are related by twofold rotational NCS, with C*α* r.m.s.d being 0.232 Å. Smaller conformational deviations between the molecules are present in loop regions 248-251 and 419-424. However, a main difference is that the C-terminal end of molecule A can only be traced to Phe448, whereas for molecule B, the entire C-terminal ending at Asn453 is visible in the maps (Figure 4D) with this region being stabilized by local interactions with the neighbouring molecule A (Figure 4C). In the case of molecule A, however, the distance between the C-terminus and molecule B is larger, and no interaction is observed (Figure 4F). In other words, molecules A and B are not identical, and symmetry operations superimposing them are imperfect. This would be the case for orthorhombic symmetry, which therefore does not hold. Pseudosymmetry on the other hand explores the A-B relations as NCS, while the twinning operator occurs from scrambling the distinction between A-B and B-A pairs in the crystal.

**Figure 4.**
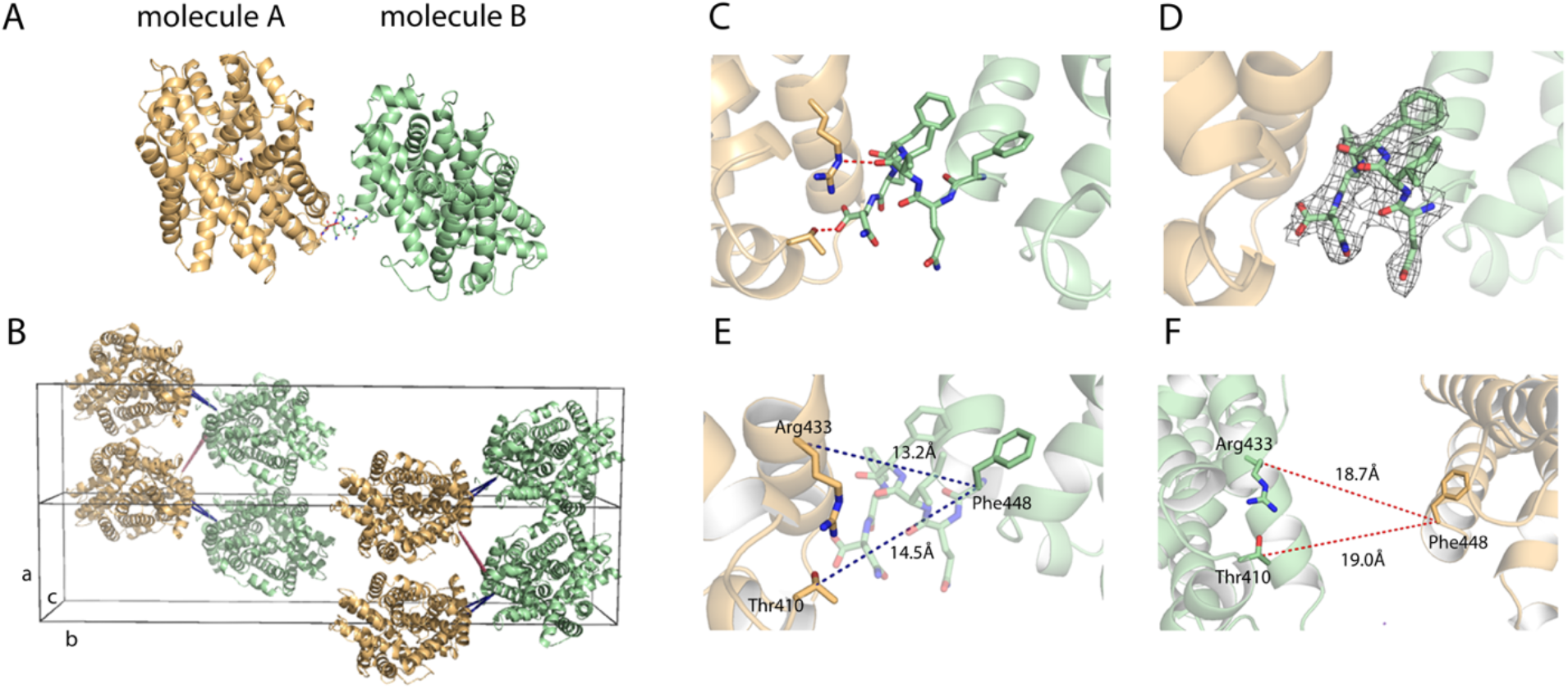
A) Two molecules in the asymmetric unit of MhsT-Val, representing also MhsT-Leu. B) Two unit cells are shown. Distances between molecules in an asymmetric unit (blue) and distances between molecules in different asymmetric units (red). C) Interactions between the C-terminus in molecule B and two residues in molecule A. D) 2F_o_-F_c_ electron density map of the additional residues in the C-terminus of molecule B contoured at 1 r.m.s.d. E) Distances between molecules in the asymmetric unit. F) Distances between molecules in different asymmetric units.

The variations in crystal packing may be due to subtle ligand-induced conformational changes of the MhsT structure, and furthermore we cannot exclude that the presence of amino acid ligands at ∼0.5 mM concentration affects the lipid-detergent phase diagram and therefore crystallization conditions. As mentioned earlier, the aromatic substrate complexes can crystallize in both P2 and P2_1_ forms, and we therefore performed a systematic crystallization approach with controlled protein:lipid (w/w) ratios for the MhsT-Trp complex (see also (Trampari, 2021)). At protein:lipid ratios of 3/0.8 and 3/1.0 MhsT-Trp crystallized mainly in the P2_1_ form, whereas at a ratio of 3:2.25 it crystallized mainly in the P2 form. These ratios are both protein and lipid batch dependent and cryo-protection procedures seem also to have an effect, but it seems clear that e.g. protein-lipid ratios affect crystal packing preferences and highlight the importance of exploring and controlling these ratios in crystal screening and optimization.

## 4. Dataset with pseudo-translation, MhsT-Ile

### 4.1. Data processing of MhsT-Ile

The datasets obtained from the MhsT-Ile crystals were processed with the XDS package (Kabsch, 2010) and CCP4 programs (Matthews, 1968, Winn *et al*., 2011) as described for the other complexes, although autoindexing failed in some cases, and for most crystals no data set could be collected due to low resolution or poor quality of the diffraction with streaky or split reflections. The crystal form was again monoclinic, but it was unclear if systematic absences were present along **b**, because this direction had been poorly sampled by the data collection runs before the onset of radiation damage and also due to anisotropic diffraction properties of thin plate crystals in the loop. Furthermore, crystals were generally not isomorphous, but two fairly isomorphous datasets were identified and merged, and despite a low completeness of ∼80%, the data were of a sufficient quality that we could distinguish space group assignments and perform structure determination and limited refinement.

Importantly, analysis of the data set in *Phenix*.*Xtriage* revealed a non-origin peak with size of 59.6% of the origin peak in the Patterson map at fractional coordinates (0.337, 0.5, -0.312), indicating translational NCS (tNCS), which together with the streaky reflections could explain the problematic indexing.

Being unable to distinguish P2 and P2_1_ in scaling without data along **b**, molecular replacement was performed in both space groups. Molecular replacement was performed in *Phenix*.*Phaser-MR* using MhsT-Trp (PDB 4US3, without TM5 and ligands) as a search model, proposing a model in P2_1_ with two molecules in the asymmetric unit (Matthew coefficient=2.52, solvent content=51.2%). However, the model exhibited negative LLG values and R-factors close to 50-60%. Additionally, the model could not explain the presence of the high non-origin peak in the Patterson map. When molecular replacement was extended to P1, searching for 4 molecules in the unit cell, different relations between the molecules were revealed. Here, as expected, two pairs of molecules could be distinguished, however surprisingly, they were related by a twofold NCS axis almost parallel to **b**. Discovering the possible relation in the asymmetric unit and guided by the non-origin peak in the Patterson map, models of molecules were created by use of *Apply NCS operators* in *Phenix* in P2_1_, with two molecules related by a NCS twofold axis parallel to **b** with the translational matrix derived from the non-origin peak in the Patterson map as only this relation would explain the coordinates of the peak. This solution not only explained the non-origin peak in the Patterson map, but also caused an immediate drop in R-factors at R_work_/R_free_ 0.313/0.347 under the initial refinement round.

The model was further refined in *Phenix*.*refine* ending with a final R_work_/R_free_ at 0.277/0.305, which was deemed acceptable considering the presence of tNCS, the low completeness of the data, and the overall lower resolution and quality of the data set. However, combined with an accurate and overall identical model from other high-resolution structures, a meaningful analysis could be obtained yielding important structural analysis and classification of the complex regarding the binding pocket, (Focht *et al*., 2021). Processing and refinement statistics are summarised in Table 4.

### 4.2. Pseudo-translation of MhsT-Ile

Translational NCS can often be detected by looking at the NCS operator, where molecules inside the asymmetric unit will be related only by translation, while orientations remain similar. However, a non-origin Patterson peak is not necessarily explained by direct relations of molecules in the same asymmetric unit only, but can also refer to relations between neighbouring asymmetric units. The NCS operator relating two molecules, A_1_ and B_1,_ in asymmetric unit 1 was analysed in *Find NCS operators* in *Phenix:*

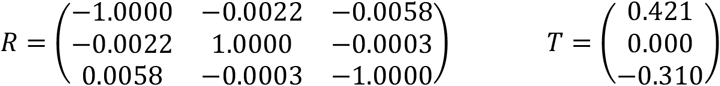

As it can be seen from the rotational matrix, the NCS operator is almost a twofold operator parallel to the crystallographic twofold screw operator. The relation between B_1_ in asymmetric unit 1 and A_2_ in asymmetric unit 2 can be calculated by the combination of the NCS operator with the twofold screw operator. In this case, the resulting operator is:

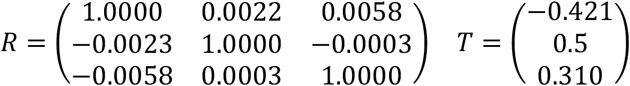

Hence, the Patterson peak corresponds to the relation between B_1_ and A_2_. The more the orientation of the related molecules deviates from each other by rotation, the smaller the off-origin peak in the Patterson map will be, and the less the tNCS will affect data processing and refinement. Normally, a deviation in orientation of the two tNCS related molecules below 10 degrees imposes a serious influence on the structure determination process (Read *et al*., 2013). Indeed, for MhsT-Ile the deviation in orientation is only 0.23 degrees (Figure 6B). This kind of tNCS can happen in cases of P2 and P2_1_ space groups with an NCS axis parallel to the crystallographic twofold or twofold screw axis, respectively. In the case, where the data would belong to the P2 space group instead of the P2_1_, the non-origin peak in the Patterson map would have the v=0.0 and not 0.5 as in the P2_1_ shown here.

Figure 5A presents the Harker section at v=0.5, whereas Figure 5B shows the stereographic projection of the self-rotation function at *k* = 180° with the crystallographic twofold screw axis and the non-crystallographic twofold axis both along **b**.

**Figure 5.**
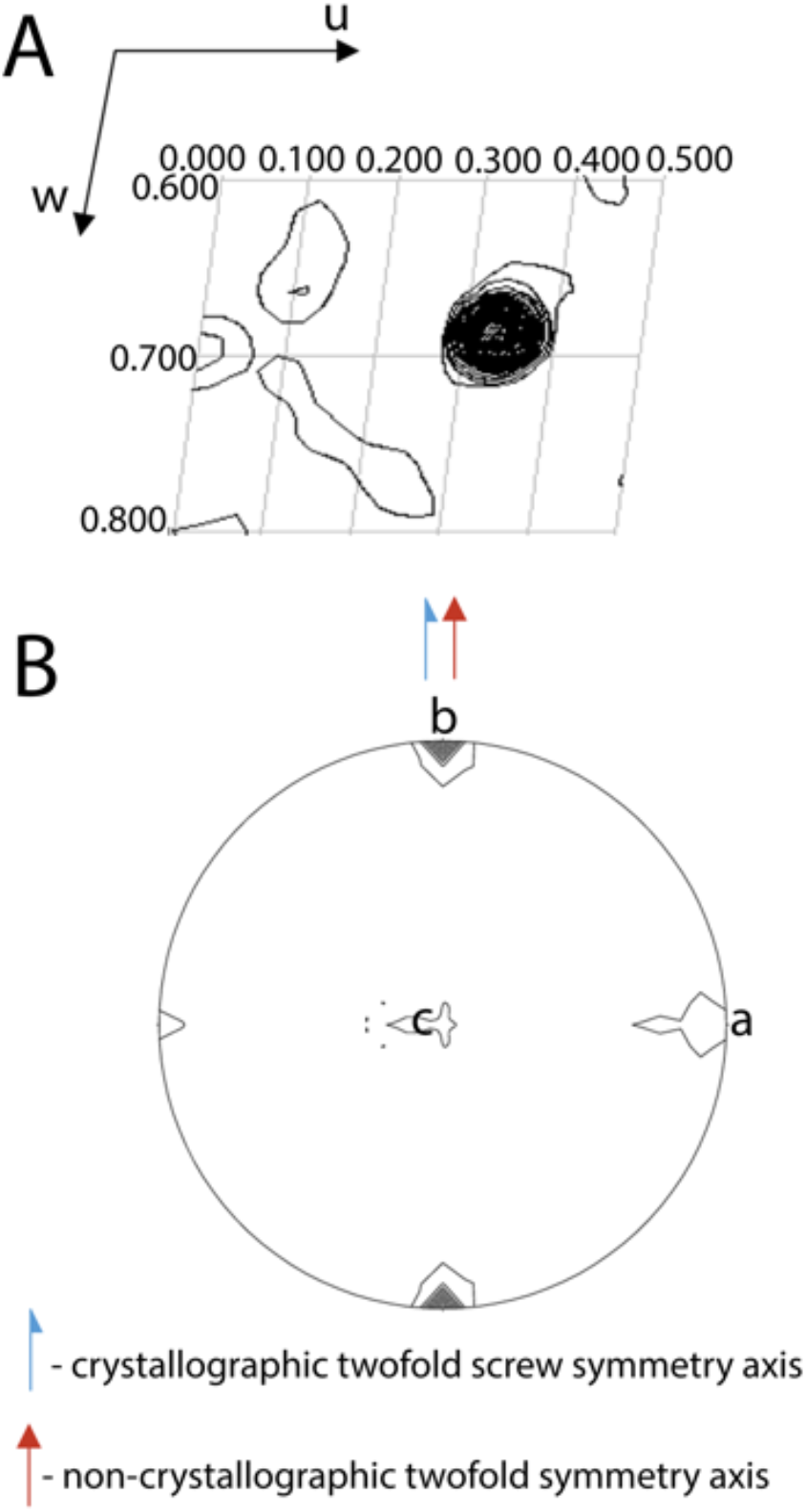
A) Harker section at v=0.5 of the Patterson map for MhsT-Ile visualizing the non-origin peak with a size of 59.6% of the origin peak. The map is drawn with a minimum contour level at 1.0 σ with 1.5 σ increments. B) Self-rotation function of MhsT-Ile. The crystallographic twofold screw axis as well as the NCS axis are both present along b.

**Figure 6.**
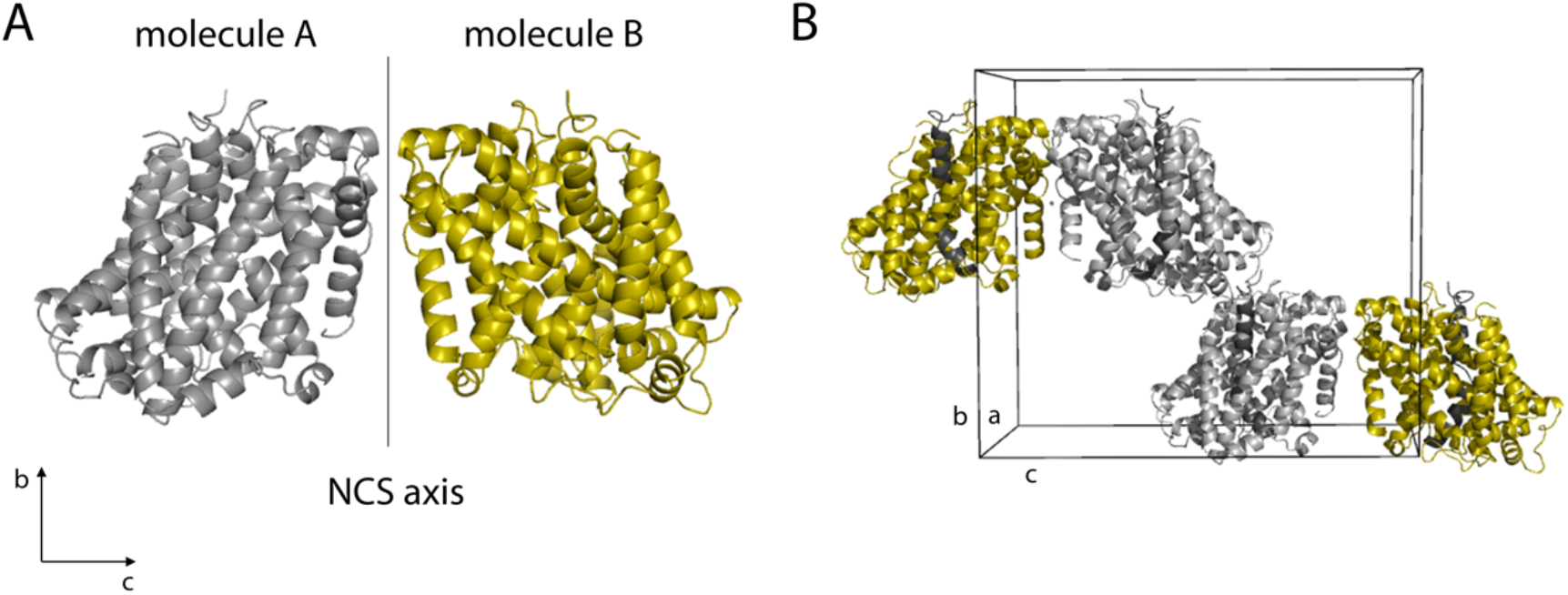
Visualization of the translational non-crystallographic symmetry of MhsT-Ile. A) Two molecules, A and B, related by NCS. The NCS axis is parallel to **b**. B) Molecule A_1_ has almost the same orientation as molecule B_2_ and these two molecules are only related by translation. TM1 is coloured in dark gray, so the orientation of the molecules can be seen easier.

## 5. Discussion

We present a remarkable case of almost identical complexes of the amino acid transporter MhsT crystallized with different amino acid substrates that exhibit a range of crystallographic phenomena including variable space group symmetries, pseudosymmetry, different degrees of pseudomerohedral twinning, and translational NCS. These variations challenged space group determination, data processing and model refinement. It is worth noting that in the two first cases (MhsT-Val and –Leu), excellent electron density maps were obtained for the pseudo-orthorhombic setting that, if combined with unreasonable presumptions of membrane protein crystal structures being allowed to pass lower quality thresholds, could lead to incorrect space group assignments and structures. In the case of twinning, at a low/absent twin fraction (MhsT-Val complex) it remains possible (in particular with hindsight) to discern the right monoclinic subgroup over orthorhombic pseudosymmetry through careful comparison of merging statistics for the individual monoclinic subgroups. However, this becomes difficult when almost perfect twinning (MhsT-Leu complex) occurs and merging statistics become essentially indistinguishable for all three monoclinic subgroups. In this case, only model refinement allowed us to distinguish the correct monoclinic space group assignment, even without twin refinement.

In the case of MhsT-Val and -Leu, the cause of pseudosymmetry can directly be identified in the crystal packing as a significant difference in local interactions around the C-terminus making two molecules A and B non-identical. The twin operation scrambles these distinctions of A-B and B-A pseudosymmetry pairs. We observe that variations in protein-lipid ratios can affect also the resulting crystal packing symmetry of the MhsT-Trp complex (also discussed in (Trampari, 2021).

Different cases of proteins determined in monoclinic P2_1_ forms with pseudomerohedral twinning have been described previously. Several reports (Larsen *et al*., 2002, Barends & Dijkstra, 2003, Golinelli-Pimpaneau, 2005) describe cases where a primitive orthorhombic symmetry is mimicked, similar to the case of MhsT. Other examples of pseudomerohedral twinning can be present in a monoclinic space group imposing apparent higher symmetry, for example when *c* · cos(β) = −a/2 (Declercq & Evrard, 2001, Rudolph *et al*., 2004), or when *a* ≈ *c* (Ban *et al*., 1999, Yang *et al*., 2000).

An analysis of entries in the Protein Database (PDB) reveals that the described types of pseudosymmetry and pseudomerohedral twinning can potentially happen quite often. Careful intensity and model refinement analyses are warranted in such cases. By considering structures determined by X-ray crystallography with experimental data available (analysis performed 20.04.2022), out of 165,026 structures in total, 27,568 (16.7%) were determined in the monoclinic P2_1_ space group. After P2_1_2_1_2_1_, it is the second-most populated space group in the PDB, and it is followed by C2. Of these, 1901 of P2_1_ structures have a *β* − angle between 89° and 91° (6.9.0%, excluding 128 entries that contain only one molecule in the asymmetric unit), where pseudomerohedral twinning could be present. Similarly, cases with model refinement stalling at suspiciously high R-factors obviously warrant careful considerations of incorrect space group assignment, pseudosymmetry and potentially twinning and translational symmetry, where also incorrect P1 assignments should be avoided.

## 6. Acknowledgements

The authors are grateful to Jesper Karlsen for advice and support of crystallographic computing

The authors are grateful to technical assistance by Tetyana Klymchuk, Lotte T. Pedersen, Anna Marie Nielsen, and support for computing by Jesper L. Karlsen.

## Notes

### Competing Interest Statement

The authors have declared no competing interest.

